# The community structure of functional brain networks exhibits scale-specific patterns of variability across individuals and time

**DOI:** 10.1101/413278

**Authors:** Richard F. Betzel, Maxwell A. Bertolero, Evan M. Gordon, Caterina Gratton, Nico U.F. Dosenbach, Danielle S. Bassett

## Abstract

The network organization of the human brain varies across individuals, changes with development and aging, and differs in disease. Discovering the major dimensions along which this variability is displayed remains a central goal of both neuroscience and clinical medicine. Such efforts can be usefully framed within the context of the brain’s modular network organization, which can be assessed quantitatively using powerful computational techniques and extended for the purposes of multi-scale analysis, dimensionality reduction, and biomarker generation. Though the concept of modularity and its utility in describing brain network organization is clear, principled methods for comparing multi-scale communities across individuals and time are surprisingly lacking. Here, we present a method that uses multi-layer networks to simultaneously discover the modular structure of many subjects at once. This method builds upon the well-known *multi-layer modularity maximization* technique, and provides a viable and principled tool for studying differences in network communities across individuals and within individuals across time. We test this method on two datasets and identify consistent patterns of inter-subject community variability, demonstrating that this variability – which would be undetectable using past approaches – is associated with measures of cognitive performance. In general, the multi-layer, multi-subject framework proposed here represents an advancement over current approaches by straighforwardly mapping community assignments across subjects and holds promise for future investigations of inter-subject community variation in clinical populations or as a result of task constraints.

## INTRODUCTION

The human brain is a complex network of functionally interconnected brain areas. Its architecture is strikingly non-random across a spectrum of scales, ranging from the local scale of individual brain areas to the global scale of the entire brain [1, 2]. Situated between these two extremes is the mesoscale comprised of sub-networks of topologically-related neural elements referred to as “communities” or modules [3, 4]. The brain’s community structure reflects regularities in its wiring diagram, delineating groups of brain areas with shared functionality [5, 6]. Critically, the brain’s community structure spans multiple organizational scales, ranging from small communities associated with functionally-specialized areas (the scale measurable with MRI) to larger communities associated with more general brain and cognitive functions [2].

Increasingly, the brain’s community structure has become the focus of many investigations. By characterizing the variability of community structure across individuals or between clinical populations, recent studies seek a deeper understanding of neuropsychiatric disorders [7, 8], development and aging [9–11], and diverse cognitive processes [12]. Despite such broad interest, there remains a paucity of principled methods for detecting and comparing communities across individuals, and little consensus on which approach maximizes advantages and minimizes disadvantages in the context of neuroscientific inquiry. Virtually all extant community detection methods rely on heuristics or make strong assumptions about the number and size of communities, the consistency of communities across individuals, and the nature of community identity as one maps community structure from one subject to another. Although these assumptions are made in order to facilitate further analyses, they can also entail unwanted biases, thereby limiting inference.

Broadly, existing approaches for comparing community structure across individuals exhibit both striking benefits and marked limitations. Consider, as an example, the popular *consensus* approach, in which group-level communities are imposed uniformly across all subjects^1^. This approach confers two notable advantages. First, the uniformity of communities permits straightforward comparisons across individuals. Second, because the communities are defined at the group level, they are likely less susceptible to overfitting. Yet, a notable disadvantage of the approach is that it precludes the possibility that communities vary across individuals. While this assumption of conservation of community structure across individuals might be reasonable in certain analyses of neurotypical cohorts (although even this assumption may be too strong; see [15]), it becomes increasingly problematic in clinical populations where heterogeneity in pathology leads to patient-specific disruptions in neuroanatomy and physiology.

Other approaches can overcome these specific issues, but are limited in different ways. Data-driven community detection methods [16, 17] naturally accommodate interindividual community variability, but can overfit noisy network data or result in situations where the mapping of communities across individuals is ambiguous or even impossible. This latter issue can be partially mitigated using template-matching techiques, such as those that register detected communities to a common set of “canonical” communities [15]. However, template-matching requires users to specify a set of template communities, restricting investigation to a single topological scale, and precluding hierarchical or multi-scale community analysis [2, 18, 19]. Moreover, template communities are oftentimes established based on resting-state data whose relationship with task-evoked community structure remains unclear [12].

Here, we propose a framework that flexibly accommodates multi-scale community analysis while unambiguously mapping communities from one subject to another. This framework builds on the well-known technique of modularity maximization [16], which algorithmically decomposes networks into internally cohesive communities and has recently been extended to be compatible with multilayer networks [20]. Past applications of multilayer modularity maximization have been largely restricted to so-called time-varying networks, where each layer represents a snapshot of a functional brain network localized to a particular window in time [21–23]. We present a modification in which network layers represent connectivity data from single subjects that are then made interdependent upon one another through the addition of inter-layer couplings.

Here, we apply multi-layer, multi-subject modularity maximization to functional connectivity data acquired as part of the Human Connectome Project (HCP) [24] and the “Midnight Scan Club” (MSC) [25]. First, we show that this approach naturally resolves ambiguities related to the mapping of communities across subjects while simultaneously recapitulating the topography of known resting-state and intrinsic connectivity networks. Next, we show that community structure varies across subjects along “modes” that are aligned with distinct organizational scales. Importantly, we show that the association of cognitive performance measures with community variability is also scale-dependent, emphasizing the necessity of detecting and analyzing community structure at multiple topological scales. Finally, using MSC data, we replicate modes of inter-subject community variability, but show that these modes differ from those associated with *intra*-subject community variability, reaffirming recent findings that variability of network organization within individuals is unique and may be a powerful source of behavioral variation. Our findings showcase the relevance of multi-scale community analysis and present methodology that will reduce the ambiguity typically associated with mapping and comparing communities across cohorts of individuals.

## RESULTS

The brain’s community structure reflects cohesive groups of functionally related brain areas and spans multiple organizational scales. inter-subject variability in the community assignments of particular brain areas has been associated with an individual’s disease, developmental, and cognitive state [7–11]. Current methods for studying this variability suffer from performance issues and tradeoffs that limit their utility. Here, we present an extension of the multi-layer modularity maximization framework to accommodate multi-subject datasets. This extension addresses many of the existing shortcomings associated with current methods and seamlessly maps communities across individuals and scales. In this section, we present the results of applying the multilayer, multi-subject modularity maximization framework to functional connectivity data taken from the Human Connectome Project (HCP) [24] and the “Midnight Scan Club” (MSC) [25].

### Detecting multi-layer, multi-subject community structure in the HCP dataset

#### Basic analysis

Characterizing inter-subject variability in community structure can provide valuable behavioral and clinical insight. Here, we examine patterns of inter-subject community variability using a novel community detection approach, which we apply to functional connectivity (FC) data made available as part of the HCP dataset. Specifically, we analyze “discovery” and “replication” cohorts each composed of *T* = 80 subjects (see **Materials and Methods** for preprocessing and cohort definition details). Our proposed community detection approach is based on a multi-layer variant [20] of the well-known modularity maximization framework [16]. In this approach, we treat subject’s connectivity matrices as “layers” that are consolidated in a unified multi-layer network, which serves as input to the community detection algorithm. By applying modularity maximization to a single multi-layer network object, we can detect communities in all layers (i.e. subjects) simultaneously. Critically, this allows us to preserve community identity across subjects. That is, two brain areas with the same community label are treated as members of the same community, irrespective of whether they correspond to different parts of the brain or appear in unique subjects. This feature of multilayer modularity maximization allows us to trivially map community assignments from one subject to another (Fig. 1).

**FIG. 1.**
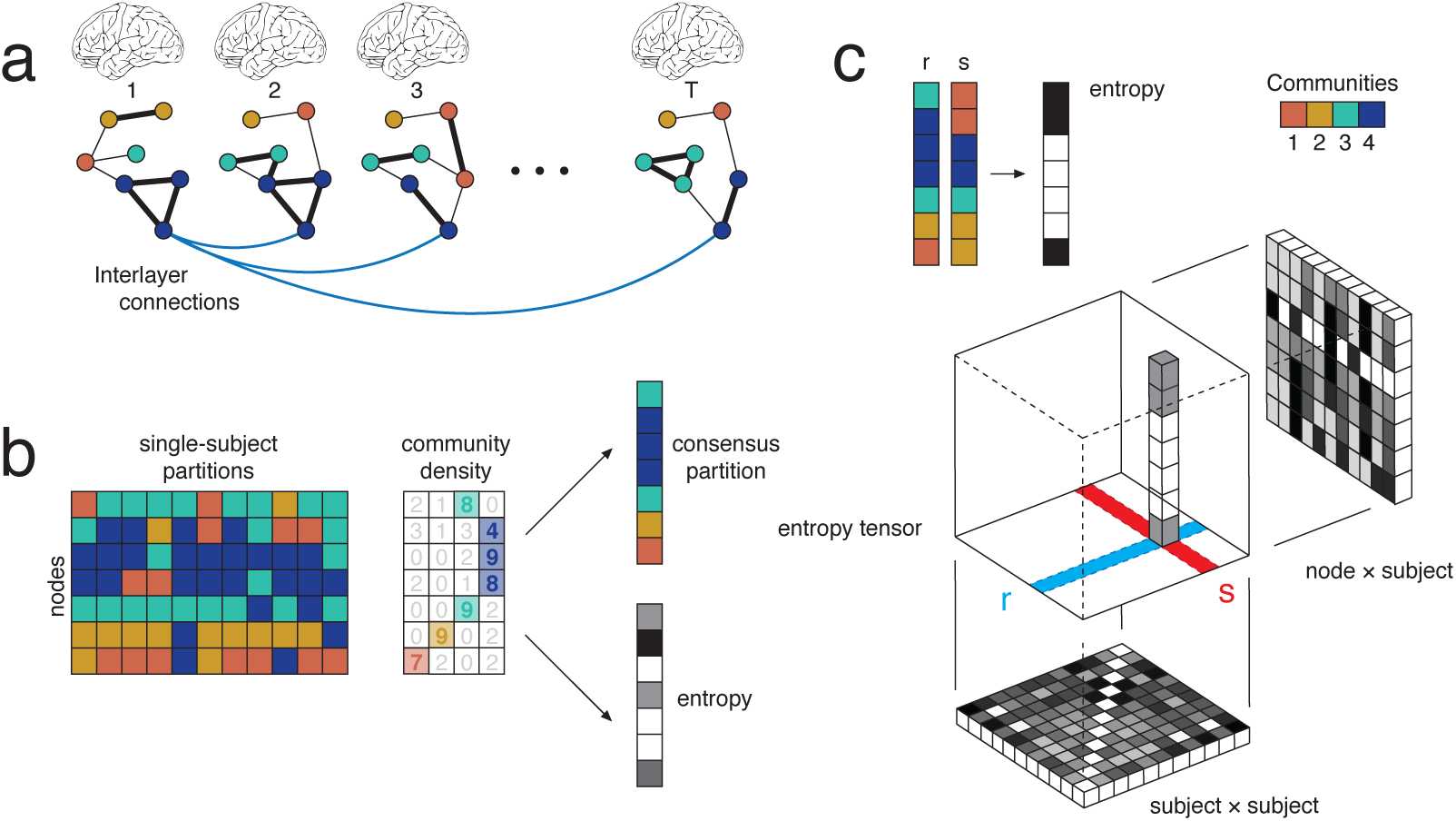
Multi-subject modularity, communities, and areal entropy. (*a*) Single-subject networks are represented as layers in a multi-layer network ensemble. Each node is linked to itself across layers, here illustrated by interlayer connections. Note that community labels are indicated by node color. (*b*) Maximizing a multi-layer modularity function returns a set of single-subject partitions. Importantly, community labels are preserved across layers; thus, if the label *C*1 appears in layers *r* and *s*, we assume that the same community has recurred. This property allows us to make several useful measurements. We can calculate, for each node, the mode of its community assignment across subjects to generate a consensus partition. We can also calculate the entropy of each node’s community assignments, which measures the variability of communities across subjects. (*c*) The preservation of community labels also allows for a direct comparison of any one subject to any other subject. Given partitions of subjects (or layers), denoted here with variables *r* and *s*, we can generate a bit vector whose values are 0, 1 depending on whether a given node has the same/different community assignment. Doing so for all pairs of subjects generates a three-dimensional *entropy tensor*. When averaged over nodes, this tensor generates a *T×T* matrix whose elements indicate, in total, the number of non-identical community assignments between pairs of subjects. When averaged over either of its other dimensions, the result is an *N × T* matrix, whose elements indicate, in total, the similarity of a node’s community assignment within a given subject to that of the remaining *T* – 1 subjects.

**FIG. 2.**
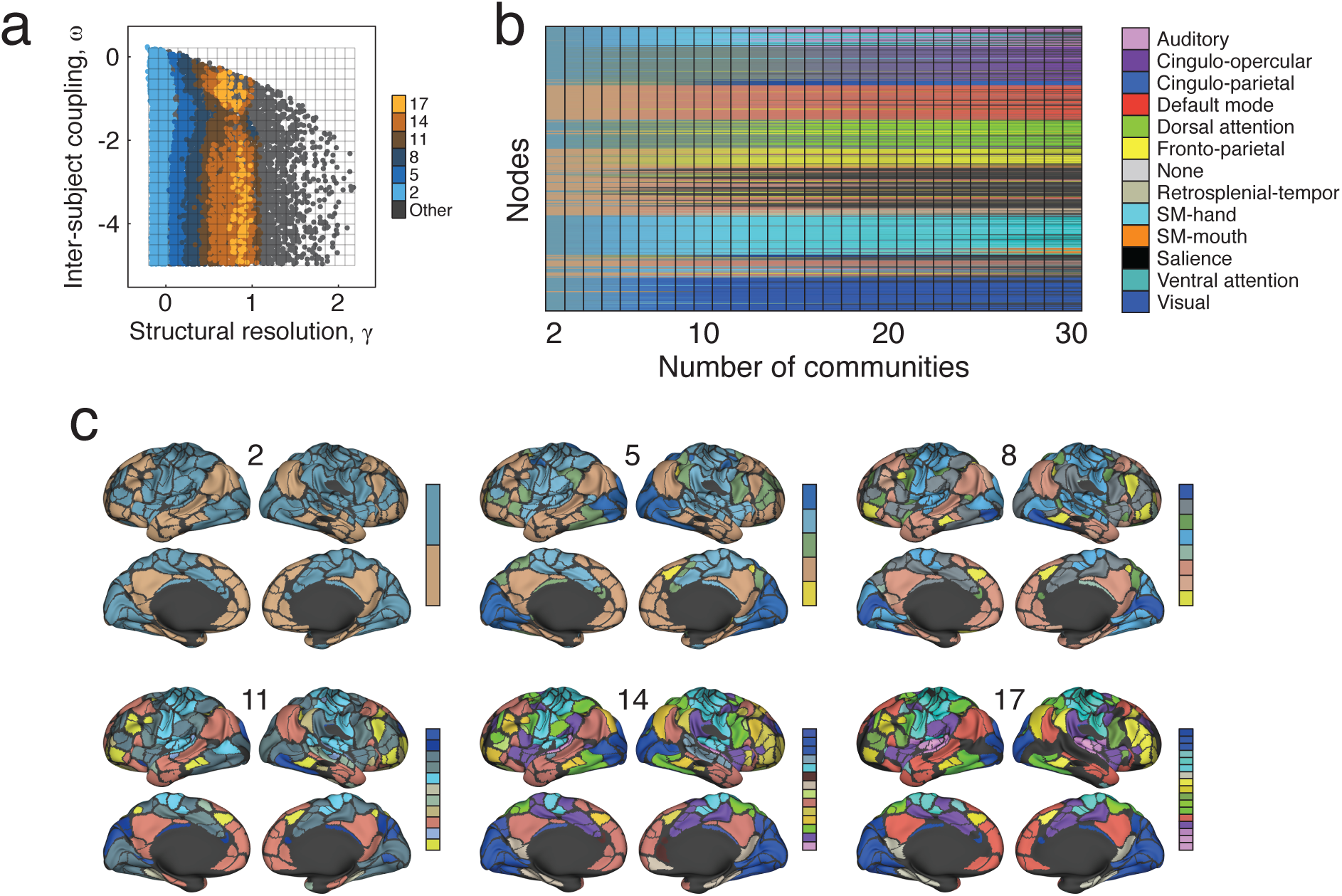
Examples of detected community structure. (*a*) The composition of detected communities depends on the structural resolution parameter, *γ*, and on the inter-subject coupling parameter, *ω*. To generate a sample of possible partitions, we chose random combinations of *γ* and *ω* and estimate consensus community structure at those points. We show, here, the locations in parameter space where the resulting consensus partitions contained 2, 5, 8, 11, 14, and 17 non-singleton communities per subject. (*b*) We ordered all consensus partitions in ascending order according to their number of non-singleton communities. Each community was colored by the weighted average of its constituent brain areas’ cognitive systems. For example, a community comprised of exclusively DMN brain areas would be assigned the DMN color (red, in this case), whereas a system composed of an equal number of DMN and visual brain areas would have a purple color (the average of the DMN’s red and the visual system’s blue). (*c*) We also show example consensus partitions as we vary the number of non-singleton communities to 2, 5, 8, 11, 14, and 17.

Multi-layer modularity maximization depends upon two free parameters. The structural resolution parameter, *γ*, determines the size of communities: smaller or larger values of *γ* result in correspondingly larger or smaller communities. The inter-subject coupling parameter, *ω*, determines the consistency of communities across layers, which in our case represent subjects: smaller or larger values of *ω* emphasize community organization that is either unique to individual subjects or shared by the entire cohort. Usually, applications using multi-layer modularity maximization focus on a restricted subset of the {*γ, ω*} parameter space, resulting in communities of characteristic size and variability. Here, however, we develop a procedure to efficiently sample a much larger region of {*γ, ω*} parameter space (see **Materials and Methods** for details). Using this procedure, we generated 40000 samples of multi-subject community structure, i.e. simultaneous estimates of each brain area’s community assignment for each subject in the cohort. We characterized each sample by measuring the variability of brain area *i*’s community assignment across subjects as the normalized entropy, *h*_*i*_. Intuitively, the value of *h*_*i*_ is equal to zero when *i* has the same community assignment across individuals and is close to 1 when *i*’s assignment is less consistent. The *N* × 1 vector *H* = {*h*_*i*_}, therefore, encodes the pattern of community variability across the entire brain. Accordingly, the average normalized entropy, 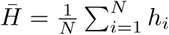, served as an index of community variability across brain areas and individuals.

As expected, we found that the number of communities varied monotonically with *γ*, the structural resolution parameter that shifts the scale at which communities are detected (Fig. 3a). Smaller values of *γ* generally resulted in larger communities while larger values of *γ* resulted in smaller communities. We also found that the number of non-singleton communities peaked at an intermediate value of *γ* ≈ 1.1 (Fig. 3b), so that the number of singleton communities increased with *γ*. In addition, we observed that the mean normalized entropy varied considerably over the full parameter space, and was greatest when both the inter-subject coupling and structural resolution parameters, *ω* and *γ*, were small (Fig. 3c).

**FIG. 3.**
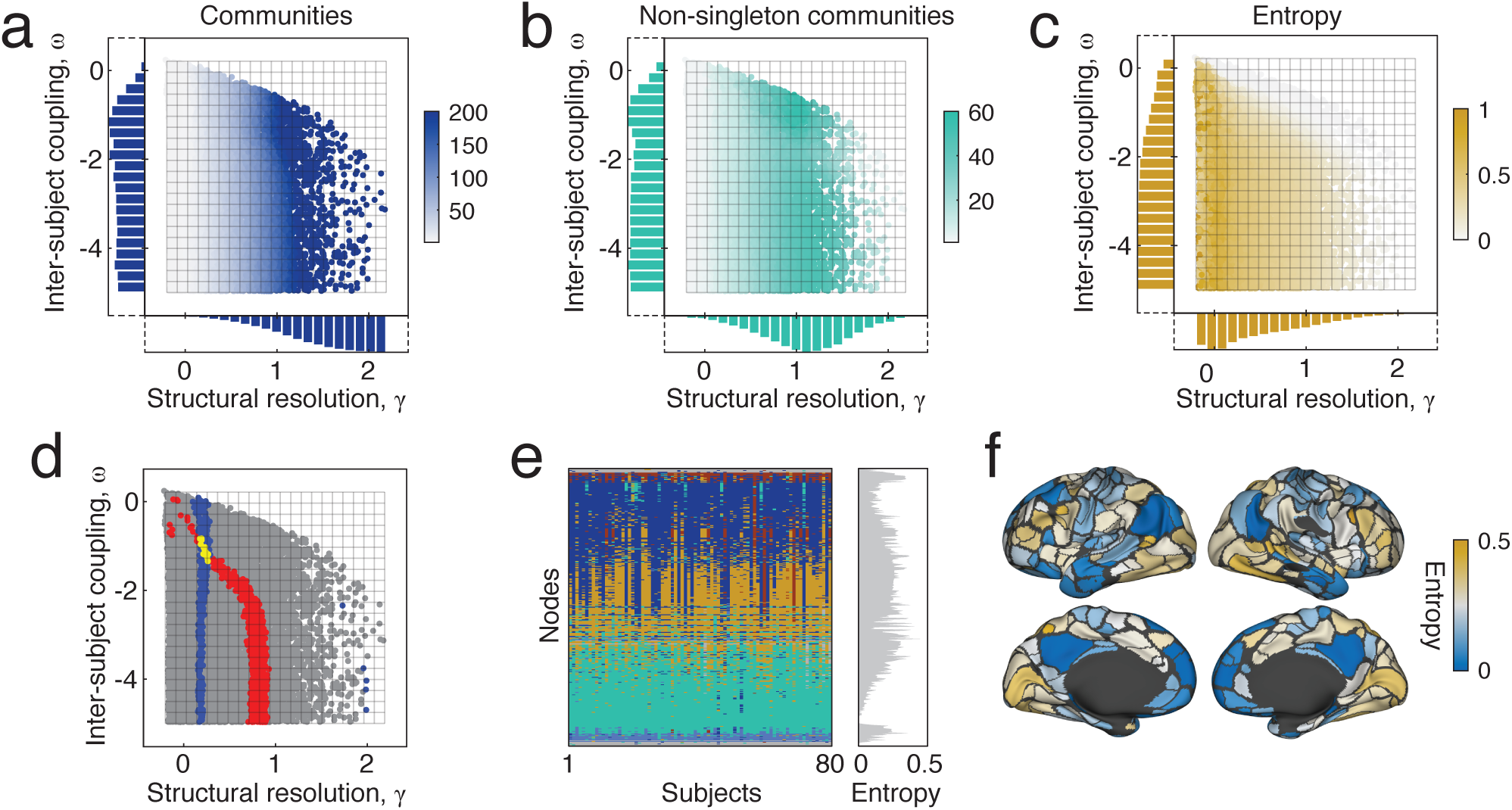
Multi-scale analysis strategy and schematic. Points are sampled in a two-dimensional constrained parameter space. The structural resolution parameter, *γ*, determines the number and size of communities while the inter-layer coupling parameter, *ω*, tunes the consistency of communities across individuals. Here, we summarize the statistics of communities detected using this sampling approach applied to the HCP333 dataset. (*a*) The number of communities per layer. (*b*) The number of communities per layer after excluding singleton communities, which are communities composed of a single node. (*c*) The mean inter-subject entropy (variability). (*d*) We can query particular subsets of partitions based on the number of communities, their average entropy, or other statistics, allowing us not only to probe different organizational scales of the network, but also to accommodate varying degrees of heterogeneity across subjects. (*e*) An example of the detected partitions and their consistency across *T* = 80 subjects. (*f*) The variability (inter-subject entropy) of community assignments across individuals. Brighter (gold) coloring indicates greater levels of variability.

Our community detection approach was designed to uncover communities of different sizes and with varying inter-subject consistency. However, some hypotheses may be easier to test by focusing on communities with a more restricted set of characteristics and statistics, such as a given number or size or a given level of inter-subject variability. Our approach naturally allows us to test these more focused hypotheses. As an example, we could specify the set of all partitions resulting in six communities (Fig. 3d; blue points) or the set of all partitions resulting in a particular level of inter-subject variability (average normalized entropy between 0.2 and 0.25) (Fig. 3d; red points). These partitions, or even their intersection (Fig. 3d; yellow points), could be extracted for additional analyses, allowing for a more detailed and nuanced exploration of community structure across subjects. As an example, we show detected communities and community entropies in Fig. 3e,f corresponding to one of the yellow points in Fig. 3d.

Collectively, these observations illustrate the mechanisms by which modularity maximization can be used to generate estimates of multi-subject community structure. Due to the two free parameters in the optimization, the method has marked utility in detecting communities at different organizational scales and with varying levels of consistency across subjects, motivating further characterization of community structure across the *γ, ω* plane.

#### Principal component analysis and modes of inter-subject variation

The sampling procedure described in the previous section generated 40000 estimates of multi-scale, multisubject community structure. For each sample, we characterized the inter-subject variability of communities as a normalized entropy vector, *H*. An important practical question is whether these patterns of variability are themselves variable, and if so, whether that variability is structured in some meaningful way. If, for example, community structure varies across subjects differently depending upon the size and the number of detected communities, such variability could have profound implications for any study of community-level correlates of behavioral measures or clinical scores. To address this possibility, we stored the full set of entropy vectors in a 333 x 40000 matrix (each column corresponds to a single sample). We column-normalized this matrix and then performed a principal component analysis (PCA), generating 332 orthonormal vectors (principal component scores) and their relative contribution to each of the 40000 entropy estimates (principal component coefficient). That is, PC scores generate brain maps while PC coefficients are defined in the parameter space. Intuitively, can be thought of as “modes” by which communities varied across subjects.

To test whether the PC scores generated here were meaningful, both practically and statistically, we performed two confirmatory analyses. First, we compared PCs calculated from discovery and replication datasets. In general, we found excellent correspondence across cohorts, with strong one-to-one PC score correlations persisting over, at least, the first twenty PCs (See Fig. S1a). Second, we compared the cumulative variance explained by PCs in the discovery cohort with the cumulative variance explained under a null model in which elements of the entropy matrix were permuted randomly and independently within columns. We found that the variance explained by the observed data was greater than that of the null model for up to thirteen components (*p* ≈ 0 based on 100 repetitions of a null model in which the elements of entropy vectors were uniformly and randomly permuted; See Fig. S1b,c). Collectively, these findings indicate that PC scores were largely replicable across two non-overlapping groups of human participants and that the total variance explained by the first few PCs exceeded that of a chance model.

Next, we characterized PC properties including their localization in parameter space and cortical topography. First, we observed that the PCs were highly localized, both in terms of their location in parameter space as well as their cortical topography. Focusing on the first four components collectively explaining *≈*78% variance, we found that PC coefficients were largely non-overlapping, tiling the parameter space and varying as a function of the structural resolution parameter, *γ* (Fig. 4a,d,g,j). This tiling phenomenon indicates that the “modes” by which communities vary across subjects are scale-dependent, varying with the number and size of detected communities.

**FIG. 4.**
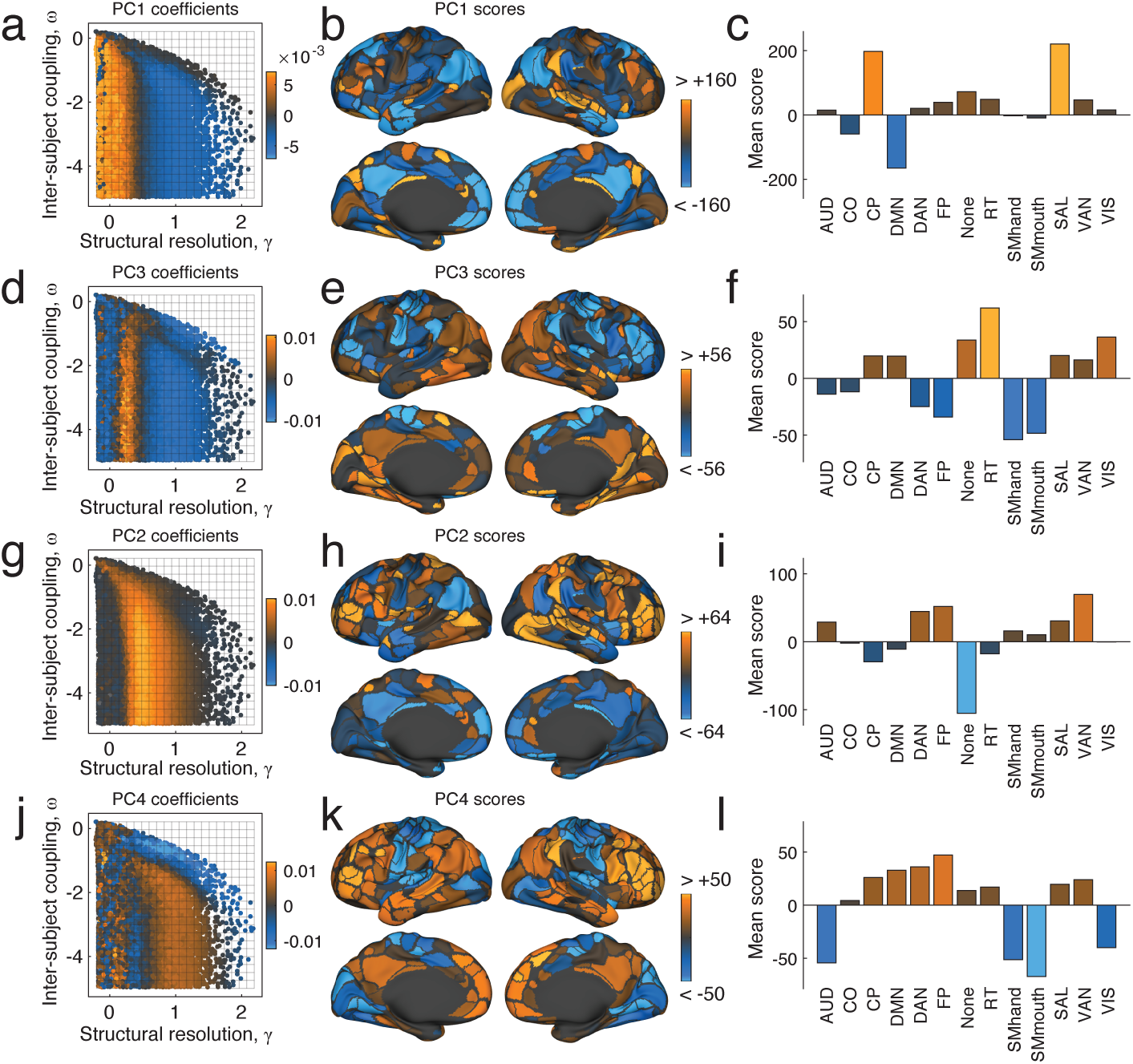
Modes of inter-subject community variability. Principal component coefficients and scores for the first four components. (*a*) PC coefficients for the first component projected into *γ, ω* parameter space. (*b*) PC scores for the same component projected onto the cortical surface. The community assignments of bright orange brain areas are highly variable across subjects at orange points in the parameter space. Conversely, the community assignments of blue brain areas are highly consistent across subjects at those same points. (*c*) Areal values of principal component scores averaged across thirteen previously-defined cognitive/functional systems [26]. This panel helps shift focus away area-level community variability and onto system level patterns of variation. The remaining panels show corresponding plots for *PC*2, *PC*3, and *PC*4. Note: we present principal components in the order in which they are expressed along the *γ* axis. This choice results in the following ordering: *PC*1, *PC*3, *PC*2, and *PC*4.

Next, we focused on the topographic distribution of PCs across the cortical surface. We found that the first component, *PC*_1_, which corresponded to relatively small values of *γ* where communities were large and few in number, implicated areas that make up the default mode (DMN), cingulo-parietal (CP), and salience systems (Fig. 4b,c). In particular, the PC scores of regions within the DMN were low (high consistency across individuals) while the PC scores of regions within the CP and SAL were high (low consistency and high variability across individuals). Across the first four principal components, different patterns of inter-subject variability were apparent. For example, in the case of *PC*_3_, we observed that the community assignments of the retrosplenialtemporal system (RT) were highly variable across individuals, while the dorsal attention (DAN), fronto-parietal (FP), and somatomotor (SMhand; SMmouth) systems were particularly stable (Fig. 4e,f). In the case of *PC*_4_, on the other hand, variability in virtually all primary sensory systems tended to be low, including auditory (AUD), somatomotor (SMhand; SMmouth), and visual (VIS), while the variability in community assignments across higher-order systems tended to be high (Fig. 4k,l).

Taken together, these findings suggest that intersubject community variability is not well-summarized by a single spatial pattern nor is it localized to a particular cognitive or functional system. Rather, the variability of community assignments across subjects depends on topological scale, as operationalized by the number and size of identified communities. This point is important, as the number and size of detected communities is usually determined by a user-defined resolution parameter [27, 28], implying that the pattern of inter-subject variability in communities can effectively be tuned by the user. This fact has important implications for applications in which one wishes to understand the relation between some aspect of network community structure and a clinical or cognitive outcome. In particular, these findings suggest that patterns of inter-subject variability can be modulated by a resolution parameter, which in principle could result in different patterns of community-behavior correlations. We explore this possibility in greater detail in the next section.

#### Brain-behavior correlations are scale dependent

One approach for relating brain network architecture to behavior involves associating measures of community structure with a behavioral, cognitive, or clinical measure of interest. In the previous section, we demonstrated that community structure varies across subjects along distinct scale-dependent modes, suggesting that brain-behavior associations could exhibit similar dependencies. This observation has important practical implications; if brain-behavior correlations are multi-scale, then any study focusing on a single organizational scale may fail to fully characterize all relevant patterns of brain-behavior correlations. To test whether this was indeed the case, we calculated a subject-level analog of the normalized entropy measure, *h*_*ir*_. This score measures for each node, *i*, and for each subject, *r*, the fraction of all other subjects in which *i*’s community assignment differed at a given point in parameter space (see **Materials and Methods**). Intuitively, this measure assesses the similarity of that node’s community assignment to its assignment in other subjects. We calculated this measure for each subject and assessed whether its variability across subjects was correlated with any of four behavioral indices related to social (SOC) or relational cognition (REL), language (LANG), or working memory (WM) task performance. Note that the derivation of these indices has been described elsewhere [29, 30] and are also briefly summarized in **Materials and Methods**.

To assess whether there was any evidence of scale-dependent patterns of brain-behavior correlations, we identified the points in parameter space where PC coefficients were greatest for the first four PCs (Fig. 5a). Separately for each PC, we computed the average correlation of inter-individual community variability with behavioral indices for each node. In general, we observed that the cortical topography of these correlation patterns varied both across behavioral indices and across the different PCs (Fig. 5b). To further illustrate this observation, we show examples of brain-wide correlation patterns of subject-level normalized entropy with the working memory (WM) behavioral index (Fig. 5c-j; the results for other behavioral indices are included in Supplementary Figs. S2 and S3). In general, the correlation pattern varied across parameter space. Depending upon where in the (*γ, ω*) parameter space the correlations were computed, the correlation magnitude was largest within different brain systems. For example, examining correlation patterns at parameter values with strong loadings onto *PC*_1_, we find evidence of system-level correlations within cingulo-opercular (CO) and visual (VIS) systems, and anti-correlations in default mode (DMN) and both dorsal/ventral attention (DAN/VAN) systems (Fig. 5e). These correlation patterns are contrasted with those observed at *PC*_4_, which are dominated by a strong positive correlation in the somatomotor (SMhand) system. Together, these findings demonstrate that not only does the brain’s modular architecture vary across individuals in a scale specific manner, but that associations of this variability with behavior also vary in a scale-specific manner. This observation motivates the re-exploration of previously analyzed data, wherein multi-scale associations might have been overlooked, and should spur the collection and analysis of datasets designed to tease apart scale-specific effects.

**FIG. 5.**
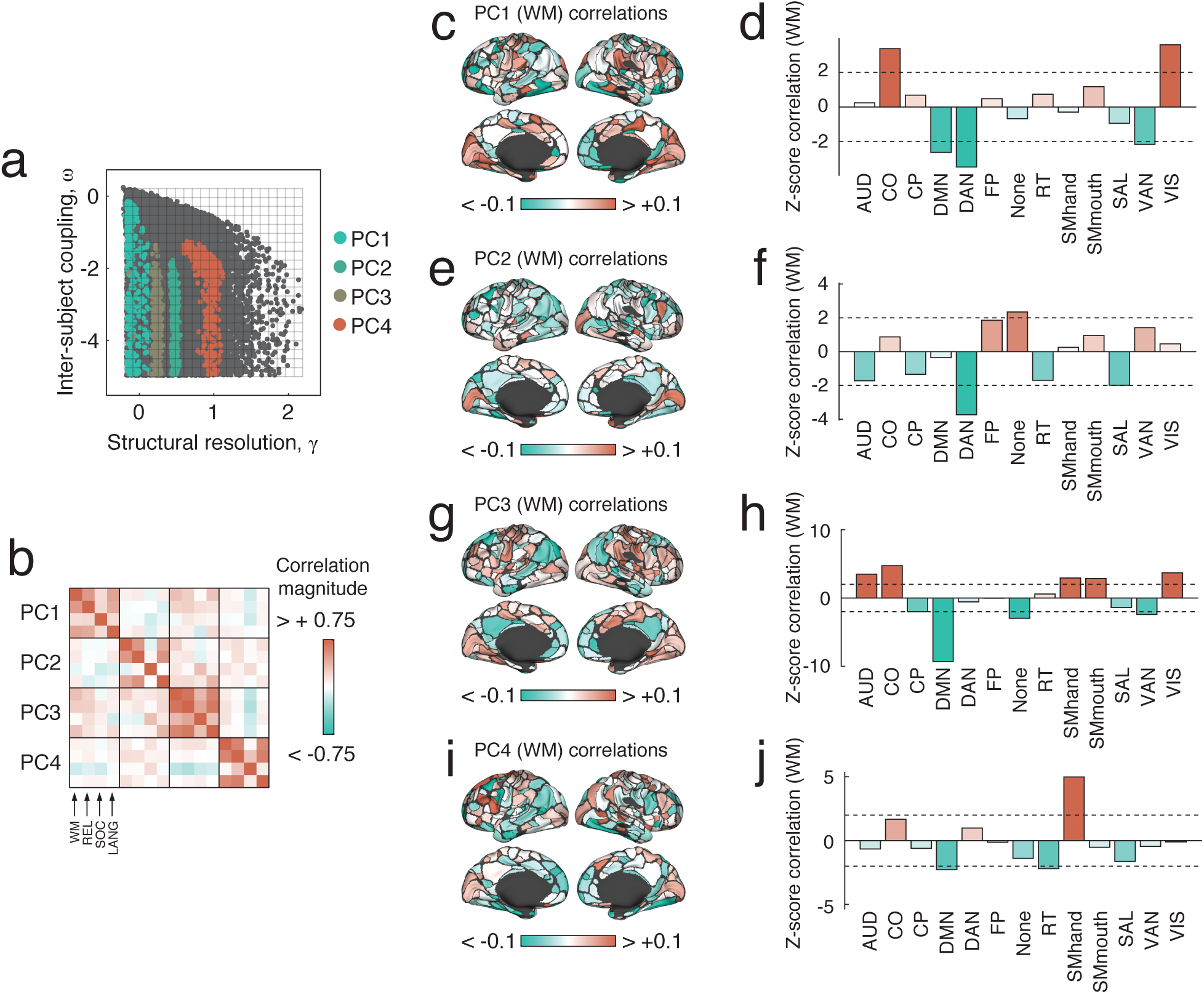
Correlation of community structure with measures of task performance. (*a*) For each PC, we studied the sub-sample of partitions corresponding to the 1% largest PC coefficients. (*b*) For each subsample, we calculated the Pearson correlation coefficient between subjects’ community entropy scores and four measures of in-scanner performance on cognitively demanding tasks: working memory (WM), relational (REL), social (SOC), and language (LANG), in the HCP terminology. In panels (*c,e,g,i*), we show the brain-behavior correlation coefficients for the first four PCs associated with performance on the WM task, plotted on the cortical surface. In panels (*d,f,h,j*), we show the mean brain-behavior correlation coefficients for the first four PCs, *z*-scored within each cognitive system. Larger *z*-scores indicate that the average correlation over all brain areas in a given system is greater than expected in random systems of the same size. Here, as an example, we show correlations for the WM task. Results for other tasks are included in the Supplementary Materials. Note that the dotted lines in panels (*d*), (*f*), (*h*), and (*j*) correspond to *z* = ±2.

### Detecting multi-layer, multi-subject community structure in the Midnight Scan Club dataset

In the previous sections we used multi-layer modularity maximization to detect and characterize patterns of inter-subject community variability in multi-subject cohorts. While network organization certainly varies across individuals, networks also vary within an individual over successive scans separated by hours, days, or weeks [25, 31–33]. Within-subject variability along different dimensions of network organization has proven useful for explaining variation in task state [34], level of attention [35], and affective state [23].

Here, we repeated our previous analyses using the recently published “Midnight Scan Club” (MSC) dataset in which ten participants underwent repeated fMRI scans (10 times per subject). Using these data we aimed to compare patterns of inter- and intra-subject variability in the modular organization of functional brain networks. To accomplish this aim, we performed two rounds of multi-layer modularity maximization. For each subject, we first generated an average connectivity matrix by aggregating usable fMRI BOLD time series data from across all resting scan sessions, and then we calculated the inter-areal correlation matrix. This procedure resulted in ten connectivity matrices (one per subject) that were treated as the layers of a multi-layer network model. As part of our second analysis, we focused on individual scan sessions from the six subjects (MSC01, MSC02, MSC03, MSC05, MSC06, and MSC07) with at least 300 low-motion frames in each of their ten scan sessions. We then estimated session-specific connectivity matrices for each subject and constructed their respective multi-layer network model whose *t*-th layer represented a subject’s connectivity pattern on the *t*-th scan session. As with the HCP dataset, we used our multi-scale sampling procedure to sample values for the structural and inter-layer coupling parameters detect communities at those points in parameter space. As before, we calculated normalized entropy measures to characterize patterns of variability across layers (either subjects or scan sessions) and to decompose these patterns into principal component scores and coefficients using SVD. By comparing the PCs generated from the inter-subject analysis with those generated by the intra-subject analysis, we could effectively identify differences in patterns of inter-*versus* intra-subject community variability.

To do so, we first assessed patterns of inter-subject variability by examining the multi-layer network where each layer corresponded to the session-averaged connectivity matrix for a different subject. Interestingly, despite the fact that these data were independently acquired and processed using a different pipeline, we found similar patterns of inter-subject variability in the MSC dataset as in the HCP dataset. We focus, in particular, on the first three PCs, which we denote MSC *PC*_1_, MSC *PC*_2_, and MSC *PC*_3_. As with HCP *PC*_1_, the brain areas in MSC *PC*_1_ associated with the lowest inter-subject variability were concentrated in the DMN (Fig. 6a). This similarity is further illustrated by computing the system-averaged correlation of MSC *PC*_1_ scores and HCP *PC*_1_ scores; we observed that the two variables exhibit a strong positive correlation (*r* = 0.79, *p* < 0.05; Fig. 6d). We find analogous relations by pairing MSC *PC*_2_ with HCP *PC*_2_ (Fig. 6b,e), and by pairing MSC *PC*_3_ with HCP *PC*_4_ (Fig. 6c,f).

**FIG. 6.**
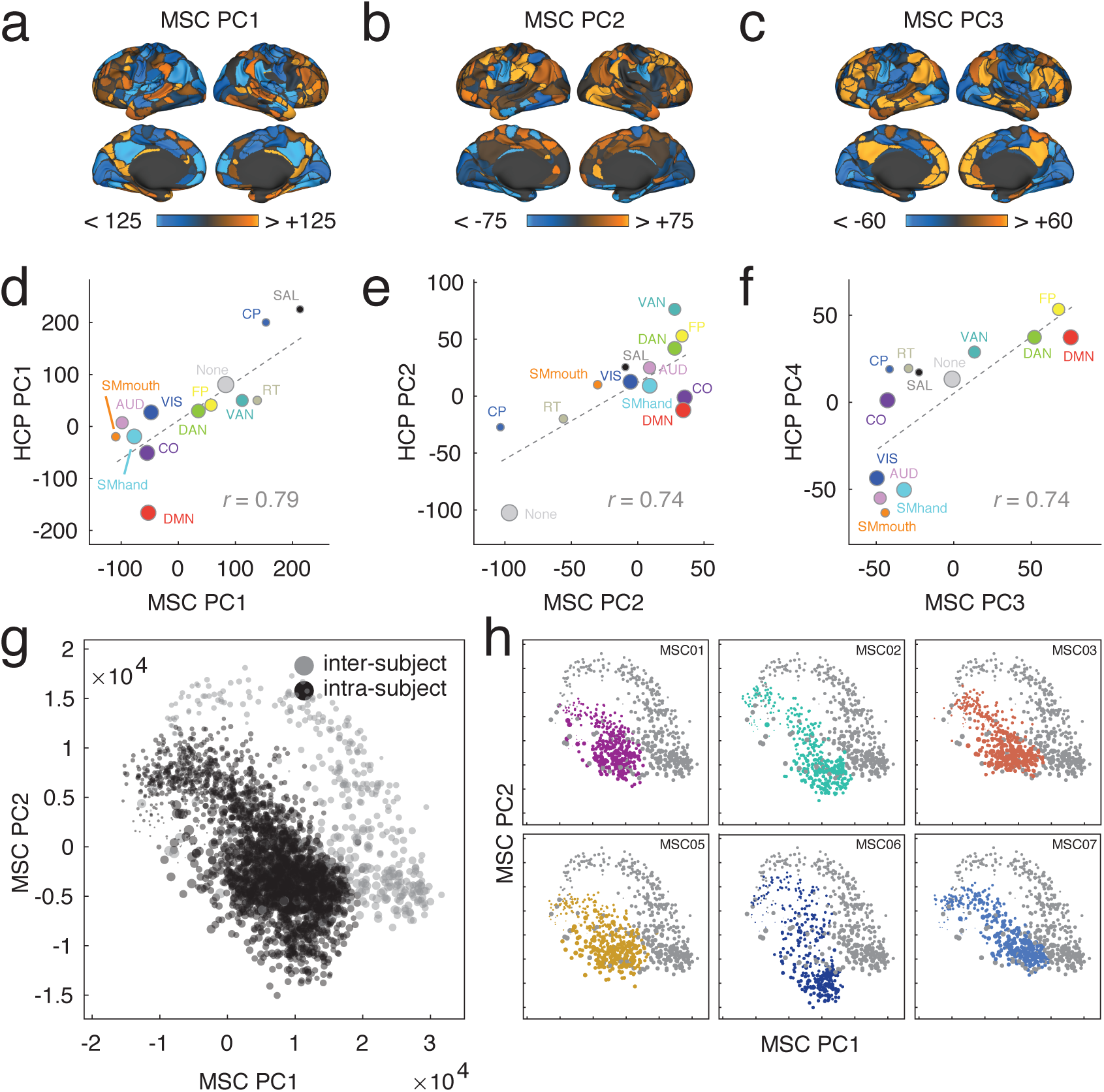
Summary and analysis of the midnight scan club dataset. In panels (*a*)-(*c*), we show surface maps depicting the first three components generated from PCA analysis of the midnight scan club (MSC) dataset. These principal components correspond, broadly, to the components detected in the HCP dataset. We compare the two datasets by averaging principal components within brain systems and computing system-average correlations. The results are shown in panels (*d*)-(*f*). The size of dots is proportional to the number of nodes assigned to each system. We perform a similar analysis of *intra*-subject variability, in which we characterize variation in community structure within subjects across scan sessions. To visualize these results, we project within-subject community entropy scores into the two-dimensional space defined by the first two intersubject principal components. In panel (*g*), we show projections of all subjects aggregated. In panel (*h*) we show subject-level projections for all six subjects individually. In panels (*g*) and (*h*), the dot size indicates the average community entropy.

Next, we performed a series of analyses to characterize *intra*-subject variability in community structure. As noted earlier, this process entailed constructing multilayer networks for the six MSC subjects with greater than 300 frames of low-motion data in each of their ten scan sessions. In these multi-layer networks, layers corresponded to functional connectivity estimated for single sessions. Otherwise, we repeated our analysis exactly as before. To compare patterns of intra- and inter-subject variability, we projected single-subject entropy patterns onto the space defined by MSC *PC*_1_ and MSC *PC*_2_. If the patterns of community variability observed within subjects were similar to those observed between subjects, then we would expect these projected intra- and inter-subject variability patterns to overlap in this lowdimensional space. However, we find that the opposite is true (Fig. 6g), and that the two categories of variability naturally separate from one another. One possible explanation is that the day-to-day variability in subject-level communities is driven by personalized factors such as sleep duration, sleep quality, affective state, or level of arousal [23, 35], whereas inter-subject community variability may be driven by temporally stable anatomical factors. In fact, we also observe subtle distinctions between the projected locations for some of the subjects (Fig. 6h). For example the bulk of points for MSC06 and MSC07 are largely non-overlapping, suggesting that not only are the modes of inter- and intra-subject variability distinct, but that intra-subject variability may exhibit some individualized patterns.

These findings consider the modes of inter-subject variability reported in the HCP data and reproduce them in an independently acquired dataset. Our results also build upon recent work characterizing intra-subject variability in functional network organization [25, 34], confirming that the patterns by which community structure varies within individuals is largely distinct from inter-subject patterns of variation. These observations suggest that the detected modes are not easily dismissed as artifacts of a particular acquisition strategy, scanner, or processing pipeline. Rather, these results suggest that communities reconfigure across individuals and *datasets* in a robust, general, and multi-scale manner [2], motivating the exploration of techniques designed to detect communities at different organizational levels.

## DISCUSSION

In this paper we adapted the popular multi-layer modularity maximization framework so that each layer represented the functional connectivity network of a single subject or scan session. This conceptual alteration facilitated unambiguous comparisons of community structure across subjects and time, permitted the straightforward calculation of consensus communities, and helped us to localize patterns of inter- and intra-subject variability. These advances made it possible for us to detect robust “modes” or (brain patterns) of inter-subject variation in community structure, which we showed were distinct from intra-subject variability. In summary, our findings offer new insight into the multi-scale community structure of functional brain networks and how that structure varies across different putative cognitive systems. The methodological advances presented here enable future studies to investigate variation in community structure across subjects from different clinical cohorts, behavioral states, and (as has already been investigated) time points.

### Advantages over current methodology

Here we propose an extension of multi-layer modularity maximization for studying how community structure varies across subjects. Our approach, which has been suggested before but never realized [36, 37], offers distinct advantages over existing methods. First, because community assignments are determined simultaneously for all subjects and because community labels are preserved across layers, we avoid the use of heuristics for mapping community assignments from one subject to another. This facilitates straightforward comparisons of community structure across individuals and allows us to easily obtain consensus communities [38]. Whereas recent work has focused on generating subject-specific community or system assignments by matching to predefined systems [15, 33] using statistical models based on shared covariance structure [39], our approach is grounded in methodology from network science, beginning with the representation of FC as a fully-weighted and signed graph, and ending with the application of modularity maximization to detect community structure [3, 13, 16].

Our use of the multi-layer modularity maximization approach for community detection is not without precedent. Past studies have leveraged this same technique to uncover the evolving community structure in timevarying FC [21–23, 40–43], across tasks [12], and motor behaviors [44]. In recasting single-subject FC matrices as individual layers and moving the problem of community detection from the level of single subjects to that of the cohort, we highlight the flexibility and generalizability of the multi-layer modularity maximization framework. It is not difficult to imagine further extensions of these methods in which network layers correspond to different connection modalities, such as structural connectivity or gene coexpression, for example [45, 46].

### Multi-scale community structure and modes of variability

In this study we characterized how communities varied across individuals and time. We found that patterns of variability could be decomposed into a series of “modes,” stereotypical and orthogonal patterns of community variability whose expression was correlated with different numbers and sizes of communities. This observation is in line with the hypothesis that brain networks, both structural and functional, exhibit community structure that spans multiple organizational scales, ranging from coarse divisions into a few large communities (e.g. the division of the resting brain into task-positive and task-negative networks [47]) to much finer divisions reflecting increased functional specialization [48]. The multi-scale and hierarchical structure of communities has a strong theoretical basis: this type of organization facilitates separation of dynamic timescales [49], efficient spatial embedding [50], evolutionary adaptability [51], and robustness to perturbations [52].

### Implications for analysis of inter-subject community variability

The observation that communities vary across subjects along distinct modes has important implications for future work. Generally, these types of studies calculate community structure at a single scale. That is, single-layer modularity maximization is performed with the structural resolution parameter set uniformly across subjects to a particular value (although this is not always the case [53, 54]). Then, variability in the detected communities is associated with some clinical score or behavioral measure. While this approach has oftentimes proven fruitful, our findings suggest that it is also limiting. Studies that focus exclusively on inter-subject variability at a single scale will fail to charactize variability in communities at other scales. Here, we demonstrated that patterns of brain-behavior correlations depend on the scale at which communities are detected. It follows, then, that any single-scale analysis of the association between community structure and clinical or behavioral measures will result in only one correlation pattern, failing to fully characterize brain-behavior associations at other scales.

Notably, our observations do not detract from these past efforts. Rather, they suggest the possibility that past studies have only scratched the surface in terms of characterizing inter-subject variability in community structure, and that there likely exists a wealth of unexplored brain-behavior associations. This notion dovetails nicely with the missions of recent data-collection and data-sharing projects in the MRI community that have made large-scale datasets and statistical maps freely available to any researcher [55–57]. These datasets, some of which have already been studied through the lens of community structure [58, 59], in principle could be reanalyzed in future studies with an increased focus on characterizing multi-scale patterns of variability.

### Limitations

Here, we extended the modularity maximization approach to be compatible with multi-subject cohorts. Though this approach has clear advantages, it also suffers from some limitations. First, as with single-layer and other multi-layer formulations of modularity maximization, the composition of detected communities depends on the free parameters, *γ* and *ω*. In our application we aimed to explore the space defined by these parameters. In fact, our findings suggest that this exploration may be necessary, as brain-behavior correlations vary across parameter space. Nonetheless, it may be advantageous in some applications to focus on a particular region of parameter space. In general, choosing the “correct” values of *γ* and *ω* is difficult, although many heuristics exist. This includes, for example, selecting the parameter values that result in consistently similar partitions [59] or identifying partitions that maximally differ from an appropriate null model [60, 61]. Finally, it is also worth noting that the modularity maximization framework could be extended much further than we did here. For instance, we fixed the values of *γ* and *ω* to be uniform across all layers. It would be interesting and potentially informative to devise heuristics for determining the values of these parameters in a meaningful and subject-specific way, allowing for finer control over the character of detected communities.

Though modularity maximization facilitates our multisubject analysis, it also serves to limit the scope of our findings. Modularity maximization implicitly assumes that a network’s community structure is uniformly assortative. This means that the communities detected using this method will be internally dense and externally sparse. While assortative communities are well-suited for segregated information processing, networks can exhibit more general classes of community structure, including core-periphery and disassortative organization [3, 62]. These communities emphasize integrative information processing and cross-community interactions. Modularity maximization, however, will fail to detect communities of this type. Other methods, such as stochastic blockmodels [63], can detect more general classes of communities. Future work could build on recent applications of blockmodeling to brain network data [64, 65] while taking advantage of multi-layer formulations to study multi-subject cohorts [66–68]. In addition, the modularity maximization framework is subject to so-called resolution limits [69, 70] that, for a given set of parameters, {*γ, ω*}, render it incapable of resolving communities below some characteristic size. This limitation, in addition to the aforementioned assumptions about partition composition, motivates the continued exploration of alternative community detection methods.

An additional limitation concerns the manner in which brain network nodes (parcels) are defined. Here, we imposed an identical parcellation across all individuals, which implicitly assumes that brain areas’ locations are consistent across subjects. Recent studies, however, have shown that areas vary in their locations across individuals [71]. Consequently, it remains a possibility that intersubject community variability could be explained by systematic differences in the locations of brain areas. Here, we use group-defined parcels as this approach remains the field standard. Future work should investigate these issues more directly, for example, by exploring the effect of subject-specific parcellations on inter-subject community variability [25, 31, 33, 34].

A final consideration concerns the scalability of the multi-layer approach. Here, we studied cohorts that included (at most) 80 subjects and connectivity data from 333 brain areas. The flattened modularity tensor associated with these data had dimensions 26640 × 26640 (333× 80). The Louvain algorithm used to maximize *Q*(*γ, ω*) was quite fast, which enabled us to sample community structure from many points in the *γ, ω* parameter space. However, for much larger matrices, corresponding to greater numbers of subjects or finer parcellations, it becomes more difficult to efficiently sample such large numbers of parameters. In such cases, one may wish to consider principled methods for constructing sparse representations of the full data matrices, or for efficiently clustering large networks [72, 73].

## CONCLUSION

Here, we studied inter-subject variability in community structure by extending the well-known modularity maximization framework to multiple subjects. We find, surprisingly, that communities vary across individuals along distinct “modes” and that these modes correspond to different organizational scales, ranging from a few large communities to many small communities. Interestingly, when we compare the variation of community structure with different measures of cognitive performance, we find that the correlation patterns are scale-dependent. This observation demonstrates that analyses that calculate brain-behavior correlations based on community structure detected at a single organizational scale may fail to uncover interesting and behaviorally relevant associations. Finally, using a second dataset, we reproduce our previously observed modes of inter-subject community variability, and also show that communities vary within subjects but along different dimensions.

## MATERIALS AND METHODS

### Datasets

We analyzed functional connectivity data from two independent datasets processed using different pipelines.

#### Human Connectome Project

We analyzed data from the Human Connectome Project (HCP), a multi-site consortia that collected extensive MRI, behavioral, and demographic data from a large cohort of subjects (*>*1000) [24]. As part of the HCP protocol, subjects underwent two separate resting state scans. All functional connectivity data analyzed in this report came from these scans and was part of the HCP S1200 release [24]. Only subjects that completed both resting-state scans were analyzed. We utilized a cortical parcellation that maximizes the similarity of functional connectivity within each parcel (*N* = 333 parcels) [26].

We preprocessed resting-state data using the following pipeline. Our analyses were based on the ICA-FIX resting-state data provided by the Human Connectome Project, which used ICA to remove nuisance and motion signals [74]. We removed the mean global signal and bandpass filtered the time series from 0.009 to 0.08 Hz. To reduce artifacts related to in-scanner head motion, frames with greater than 0.2 millimeters of framewise displacement or a derivative root mean square above 75 were removed [75]. Subjects whose scans resulted in fewer than 50% of the total frames left were not analyzed further; a total of 827 subjects met this criteria for all resting-state scans.

For all scans, the MSMAII registration was used, and the mean time series of vertices on the cortical surface (fsL32K) in each of the *N* = 333 parcels was calculated [26]. We used this particular parcellation as it has high functional connectivity homogeneity within each parcel and the number of nodes is consistent with other analyses. The functional connectivity matrix for each subject was calculated as the pairwise Pearson correlation coefficient (subsequently Fisher *z*-transformed) between times series of all nodes. Both left-right and right-left phase encoding directions scans were used, and the mean functional connectivity matrix across the four resting-state scans was calculated.

In-scanner head motion is known to induce spurious correlations in resting state FC [75]. To reduce the impact of head motion on detected communities, we focused our analyses onto smaller, more exclusive subsets of subjects. Specifically, we divided the full HCP dataset into smaller discovery and validation datasets comprising 80 subjects each. We chose this number of subjects to ensure that dataset size was comparable to what is reported in the typical fMRI study. The test cohort included the 1st, 3rd, 5th … and 159th subjects, in ascending order of mean framewise displacement. The validation cohort was defined as the 2nd, 4th, 6th, … 160th subjects, ordered according to the same criterion. Note, because this procedure generated datasets with low head motion, it is possible that related subjects appear together in the same cohort.

We also analyzed HCP behavioral data. For the Working Memory task, our measure of task performance was given by the mean accuracy across all conditions (WM Task Acc). For the Relational task, our measure of task performance was given by the mean accuracy across all conditions (Relational Task Acc). For the Language task, our measure of task performance was given by the highest level reached in either the Language or Math conditions (maximum of Language Task Story Avg Difficulty Level and Language Task Math Avg Difficulty Level). For the Social task, our measure of task performance was given by the mean across the random and theory of mind conditions (mean of Social Task TOM Perc TOM and Social Task Random Perc Random).

#### Midnight Scan Club

Data were collected from ten healthy, right-handed, young adult subjects (5 females; age: 24-34). One of the subjects is an author (NUFD), and the remaining subjects were recruited from the Washington University community. Informed consent was obtained from all participants. The study was approved by the Washington University School of Medicine Human Studies Committee and Institutional Review Board. This dataset was previously reported in [15, 34] and is publicly available at https://openneuro.org/datasets/ds000224/versions/00002. Imaging for each subject was performed on a Siemens TRIO 3T MRI scanner over the course of 12 sessions conducted on separate days, each beginning at midnight. In total, four T1-weighted images, four T2-weighted images, and 5 hours of restingstate BOLD fMRI were collected from each subject. For further details regarding data acquisition parameters, see [15].

MRI data were preprocessed and sampled to the surface as described in [15] and with shared code available at https://github.com/MidnightScanClub. The steps are summarized briefly below. The high-resolution structural MRI data were averaged together, and the average T1 images were used to generate hand-edited cortical surfaces using Freesurfer [76]. The resulting surfaces were registered into fs LR 32k surface space as described in [74]. Separately, an average native T1-to-Talaraich [77] volumetric atlas transform was calculated. That transform was applied to the fs LR 32k surfaces to put them into Talaraich volumetric space.

All fMRI data first underwent pre-processing (in the volume) to correct for artifacts and align data, including slice-timing correction, frame-to-frame alignment to correct for motion, and intensity normalization to mode 1000. Functional data were then registered to the T2 image, which was registered to the high-resolution T1 anatomical image, which in turn had been previously registered to the template space. Finally, functional data underwent distortion correction [15]. Registration, atlas transformation, resampling to 3 mm isotropic resolution, and distortion correction were all combined and applied in a single transformation step [78]. Subsequent steps were all completed on the atlas transformed and resampled data.

Processing steps specific to functional connectivity were undertaken to reduce the influence of artifacts on functional network data. These steps are described in detail in [79], and include (1) demeaning and de-trending of the data, (2) nuisance regression of signals from white matter, cerebrospinal fluid, and the global signal, (3) removal of high motion frames (with framewise displacement (FD) *>* 0.2 mm; see [15]) and their interpolation using power-spectral matched data, and (4) bandpass filtering (0.009 Hz to 0.08 Hz). After this volumetric pre-processing, functional data were sampled to the cortical surface and combined with volumetric subcortical and cerebellar data into the CIFTI data format using the Connectome Workbench [80]. Finally, data were smoothed (Gaussian kernel, *σ* = 2.55 mm) with 2-D geodesic smoothing on the surface and 3-D Euclidean smoothing for subcortical volumetric data.

### Modularity maximization

A critical step in any modularity analysis is to determine the community assignments of brain areas. Because real-world networks are too complex to identify communities from visual inspection alone, their nodes’ community assignments must be determined algorithmically through a process known as community detection [13]. There are many different ways to define a network’s communities and equally many algorithms for detecting them. Of these, modularity maximization is likely the most popular [16].

Intuitively, modularity maximization operates according to a simple principle: communities correspond to groups of nodes that are more strongly connected to one another than would be expected by chance alone. The detection of these communities, in practice, is accomplished by optimizing a modularity objective function:

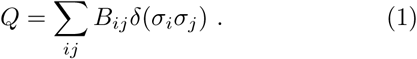

In this expression, *B*_*ij*_ = *W*_*ij*_ *–P*_*ij*_, where *W*_*ij*_ and *P*_*ij*_ are the observed and expected weight of the connection between nodes *i* and *j*. The full matrix, *B* = {*B*_*ij*_}, is referred to as the modularity matrix whose elements encode the difference between observed and expected connection weights. The variable *σ*_*i*_ *ϵ {*1, *…, K*} indicates to which of the *K* communities node *i* is assigned. The Kronecker delta function, *δ*(*x, y*), takes on a value of 1 when *x* = *y* and 0 when *x*≠*y* Finally, *Q* is the modularity quality function to be maximized. Effectively, *Q* is the sum over within-community elements of the modularity matrix. In general, larger values of *Q* are taken to indicate higher quality partitions and correspond to internally dense and externally sparse communities.

The modularity function suffers from a “resolution limit,” meaning that communities below a characteristic scale are undetectable [70]. To circumvent this issue, some applications introduce a tunable structural resolution parameter, *γ*, which scales the relative importance of the null connectivity model:

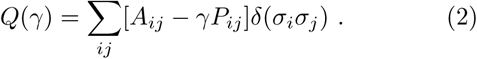

The effect of this parameterization is that optimizing *Q*(*γ*) for large values of *γ* results in the detection of many small communities, whereas optimizing *Q*(*γ*) for small values of *γ* results in a few large communities.

The modularity function has been expanded further so that it is compatible with multi-layer networks composed of distinct layers corresponding to different connection modalities (e.g., structural and functional connectivity) or estimates of network structure at different time points [20]. In the multi-layer analog of the modularity function, nodes are linked to themselves across layers by an interlayer coupling parameter, *ω*. This parameter determines the similarity of communities detected across layers, with larger values of *ω* resulting in increased homogeneity of communities across layers. The multi-layer expression for modularity is given by:

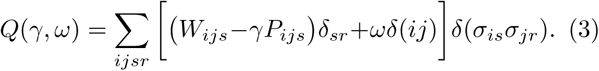

The multi-layer, multi-scale modularity, *Q*(*γ, ω*), operates on the same principle as the single-layer version. Communities are defined by placing stronger-than-expected connections within communities. The main difference between the single- and multi-layer modularity functions is that, by adding interlayer connections of weight *ω*, it can be advantageous from the perspective of optimizing *Q*(*γ, ω*) to assign nodes in different layers to the same community.

### Multi-layer, multi-subject modularity

Our approach builds directly upon the canonical multilayer modularity framework. It differs in two important ways, the first is conceptual while the second is methodological. Conceptually, rather than letting layers represent different connection modalities [45, 46], frequency bands [81, 82], or network estimates at different time points [21], we let layers represent functional connectivity matrices corresponding to single subjects. Effectively, this choice allows us to obtain estimates of community assignments simultaneously for all subjects. Importantly, this approach also preserves community labels across subjects (provided *ω* > 0), making the process of comparing communities across subjects trivially easy. Methodologically, our approach differs from the general case in that we let *P*_*ijs*_ = 1 for all *i, j, s*. This choice is in line with best practices for modularity maximization when the networks correspond to correlation matrices [83–85]. In our past work we have shown that this decision results in communities with broadly recognizable topographic features [86, 87].

As with the canonical version of multi-layer modularity maximization, the multi-subject version depends upon two free parameters, *γ* and *ω*. The value of *γ* determines the scale of detected communities, as reflected in their number and size. As before, smaller values of *γ* result in a few large communities while larger values result in many small communities. Similarly, the value of *ω* emphasizes the consistency of communities across individuals. Larger values of *ω* emphasize network features and communities that are common across subjects, whereas small values of *ω* emphasize the uniqueness of individuals’ community structures. The parameter space defined by *γ* and *ω* contains partitions of the network into communities ranging from single nodes to the whole network, that are entirely unique to individuals or perfectly consistent across individuals. Note, that while *ω* influences the consistency of communities across subjects, it is implemented globally and its influence is exerted over all subjects equally.

#### Sampling multi-scale, multi-consistency community structure

Though many studies fix the values of *γ* and *ω* in order to focus on partitions of networks into roughly the same number of communities across subjects, we aimed to explore the full range of partitions, both in terms of community size but also in terms of the consistency of community structure across individuals. In general, however, the relevant ranges of *γ* and *ω* are unknown ahead of time. In our application, we are interested in characterizing variability in community structure across individuals, and therefore wish to avoid extreme partitions; that is, combinations of *γ* and *ω* that result in:

1. singleton communities (each node is its own community),
2. whole-network partitions (all nodes are assigned to the same community),
3. partitions that are identical across all subjects, or
4. partitions that are maximally dissimilar across all subjects.

To sample community structure, we employed a novel two-stage procedure. The first stage allowed us to bipartition the parameter space defined by *γ* and *ω* into two sub-spaces, one where the above-defined criteria were true and another where the above-defined criteria were false. In the second stage we sampled parameter pairs from within this sub-space to generate a distribution of possible partitions that spanned all organizational scales and levels of consistency.

The first stage is initialized with the user defining the boundaries of a rectangular parameter space, forcing *γ* and *ω* to fall within [*γ*_*min*_, *γ*_*max*_] and [*ω*_*min*_, *ω*_*max*_], respectively. Here, we set *γ*_*min*_ = min_*ijs*_ *W*_*ijs*_, *γ*_*max*_ = max_*ijs*_ *W*_*ijs*_, *ω*_*min*_ = 0 and *ω*_*max*_ = 1. Next, we sampled parameter pairs uniformly and at random from within this sub-space. To identify regions of this sub-space in which the above-defined four criteria hold, we optimized *Q*(*γ, ω*) for each pair of parameter samples using a generalization of the so-called Louvain algorithm [88, 89]. This optimization resulted in a multi-layer partition for which we computed the mean number of communities per layer and the mean consistency of communities across layers (see the next section). Intuitively, if the average number of communities was fewer than 2 or greater than *N*, or the mean consistency of communities was equal to 0 (maximal inconsistency) or 1 (maximal consistency), we considered the corresponding point in parameter space to be “bad,”, as it would have failed to satisfy at least one of the four criteria. Parameter values that satisfy these criteria, on the other hand, are considered “good”.

Next, we calculate the local homogeneity of parameter space by calculating the entropy of each sampled point’s 25 nearest neighbors. Points with non-zero values of entropy exhibit inhomoegeneities and correspond to regions of parameter space where at least one of the four criteria is sometimes not satisfied. We identified all such points and constructed around them the unique non-convex polygon. Intuitively, partitions corresponding to parameter pairs that fall within this polygon are likely to satisfy all four criteria; points located outside this polygon are unlikely to satisfy all four criteria. We repeated this procedure five times (a total of 5000 samples), each time refining our definition of the boundary between “good” and “bad” regions of parameter space.

At this point our procedure moved onto the second stage. In this stage, we sampled 40000 partitions from within the “good” region. Values of *γ* were sampled uniformly, while values of *ω* were sampled from an exponential distribution, so that small values of *ω* are sampled more frequently than large values. For each pair of parameter values we optimized *Q*(*γ, ω*), resulting in 40000 multi-layer partitions. These partitions formed the basis for all subsequent analyses.

### Consensus communities and community entropy

Optimizing the multi-layer, multi-scale modularity expression returns *σ* = *σ*_*is*_, whose elements encode the community assignment of node *i* in subject *s*. From this two-dimensional matrix we can calculate several useful statistics that would not have been accessible using the traditional single-layer approach. First, we can obtain a consensus partition, or a group-representative set of communities. Whereas past studies have relied on iterative clustering with an arbitrary threshold to obtain consensus communities [38], the multi-layer ensemble makes their estimation straightforward. The consensus assignment of node *i* is given as:

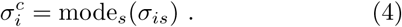

Next, we can also calculate the variability of community assignments across subjects for a given node. Let *p*_*i*_(*k*) equal the fraction of all subjects whose node *i* is assigned to community *k*. The variability of this node’s assignment can be characterized with the entropy:

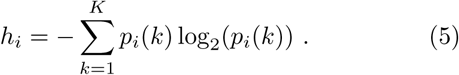

We can standardize this entropy to the range [0, 1] by dividing by log_2_(*K*).

### Principal component analysis

The entropy vector describes the variability of each brain areas’ community assignment across subjects. These vectors, in general, may depend on where in the *γ, ω* parameter space those communities were sampled. We wished to assess whether certain patterns of inter-individual community variability appeared more frequently than others and whether these patterns were localized to specific regions of parameter space.

To address this question, we subjected the column-normalized (zero mean, unit variance) matrix of entropy scores to a principal component analysis via singular value decomposition (SVD). Let *H* ∈ [*N × N*_*reps*_] be the matrix of nodes’ entropy scores across all repetitions of the Louvain algorithm. In the case of HCP and MSC data with the Gordon atlas, this matrix has dimensions [333 × 40000]. Singular value decomposition factorizes *H* into left and right singular vectors, *U* ∈ [*N × N*] and *V* ∈ [*N*_*reps*_ *× N*] and a set of singular values, σ ∈ [*N × N*_*reps*_], satisfying the relationship:

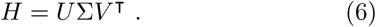

The principal component scores and coefficients are given by the columns of *U*, which are by definition orthogonal to one another, and the rows of *V*, respectively. The squared diagonal elements of Σ give the percent variance accounted for by principal components. We interpret the columns of *U* as modes of inter-individual community variation, or dominant brain-wide patterns of entropy scores.

In the case of the MSC dataset, we performed SVD on the inter-individual entropy matrix, which was calculated given communities estimated using a multi-layer network whose layers represented session-averaged FC for each of the 10 subjects. We then extracted the first three principal component scores, *U*_inter_ = [*U*_(:,1)_*U*_(:,2)_, *U*_(:,3)_], which defined a low-dimensional space of inter-subject community variability.

In addition to calculating inter-individual communities, we also applied modularity maximization to six different multi-layer networks where each layer represented FC for a given subject on a given scan session. Analogously to our calculation of inter-subject community variability, we computed inter-session community variability for each subject, and subsequently calculated each subject’s matrix of column normalized entropy scores, *H*_subject_. To project these inter-session data into the inter-subject space, we performed the multiplication: *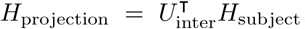*The result, *H*projection *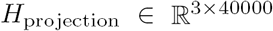* is a projection of inter-session variability patterns onto axes defined according to inter-subject community variability.

## DATA AVAILABILITY STATEMENT

Data were provided in part by the Human Connectome Project, WU-Minn Consortium (Principal Investigators: David Van Essen and Kamil Ugurbil; 1U54MH091657) funded by the 16 NIH Institutes and Centers that support the NIH Blueprint for Neuroscience Research; and by the McDonnell Center for Systems Neuroscience at Washington University.

## AUTHOR CONTRIBUTIONS

This study was designed, carried out, and written by RFB. DSB secured funding for the work. MAB processed and provided HCP MRI and behavioral data. EMG, CG, NUFD processed and provided MSC MRI data. All authors contributed to the direction of the research and edited the paper.

## ACKNOWLEDGEMENTS

RFB, MB, and DSB would like to acknowledge support from the John D. and Catherine T. MacArthur Foundation, the Alfred P. Sloan Foundation, the ISI Foundation, the Paul Allen Foundation, the Army Research Laboratory (W911NF-10-2-0022), the Army Research Office (Bassett-W911NF-14-1-0679, Grafton-W911NF16-1-0474, DCISTW911NF-17-2-0181), the Office of Naval Research, the National Institute of Mental Health (2-R01-DC-009209-11, R01-MH112847, R01-MH107235, R21-M MH-106799), the National Institute of Child Health and Human Development (1R01HD086888-01), National Institute of Neurological Disorders and Stroke (R01 NS099348), and the National Science Foundation (BCS-1441502, BCS-1430087, NSF PHY-1554488 and BCS-1631550). The content is solely the responsibility of the authors and does not necessarily represent the official views of any of the funding agencies.

---

**FIG. S1.**
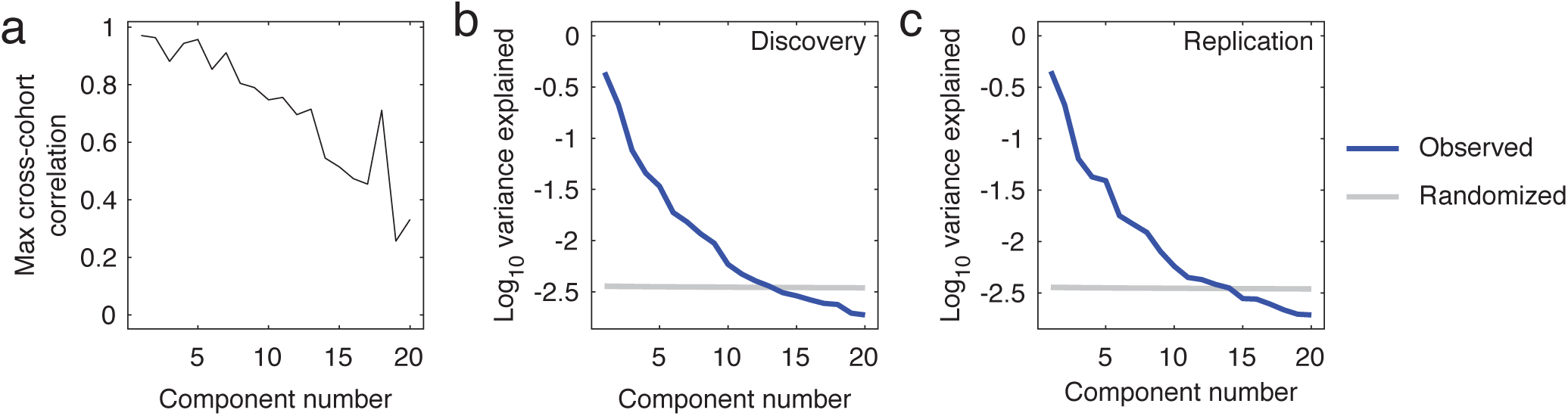
Cross-cohort analysis of principal components. (*a*) For each PC in the discovery dataset (cohort 1), we calculated the maximum correlation with any of the first 20 PCs in the replication dataset (cohort 2).(*b*) Variance explained by the first 20 principal components in the discovery dataset. The blue curve shows observed values while the gray curve shows values expected under a chance model. (*c*) Same plot as panel *b* but for the replication dataset.

**FIG. S2.**
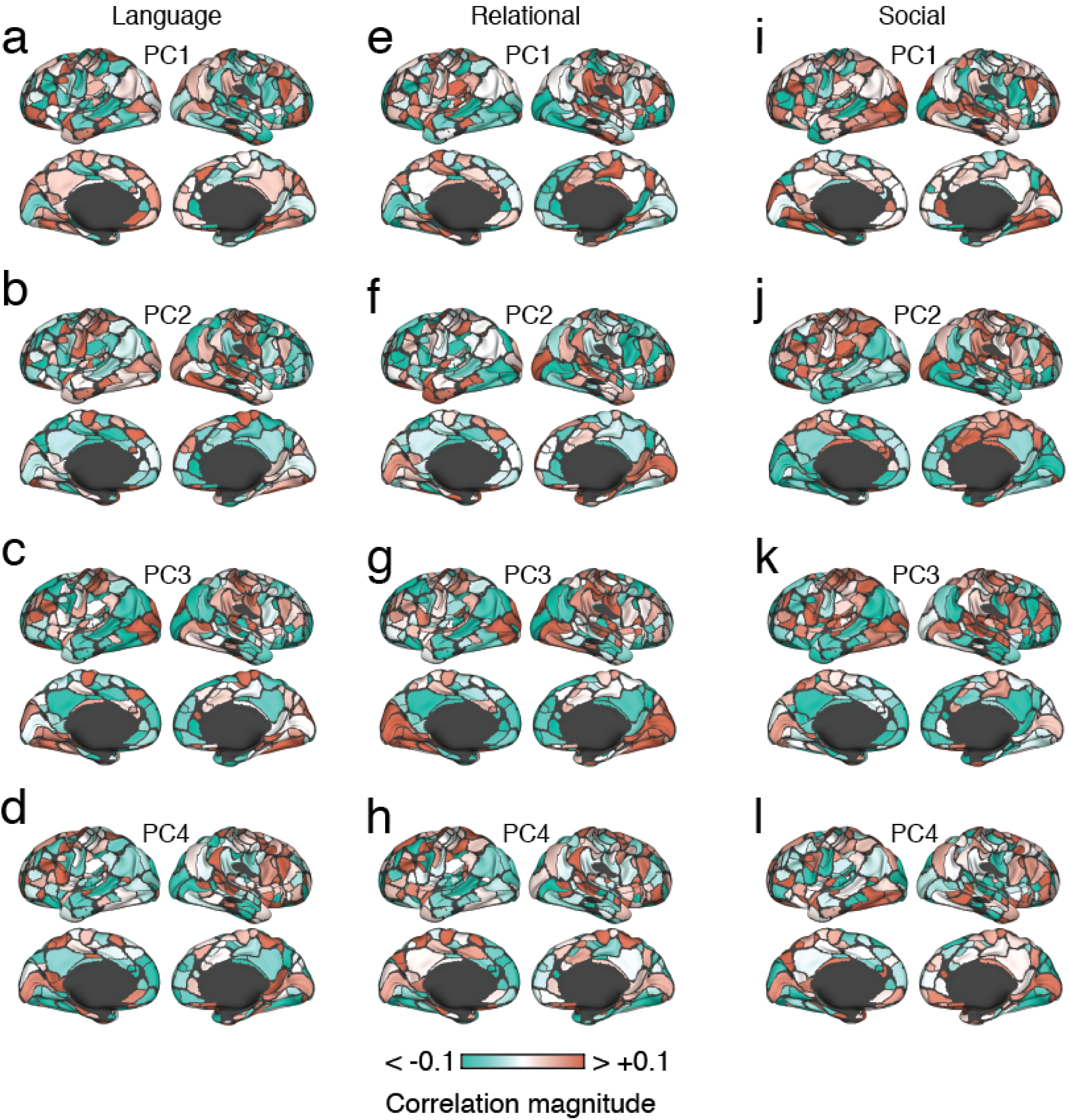
Spatial maps depicting the correlation of community structure with measures of task performance. In the main text we calculated the correlation of community variability with performance on the working memory task and plotted the strength of correlation onto the cortical surface. Here, we show similar surface plots for the Language, Relational, and Social tasks. We also computed correlations separately for the first four modes (principal components) of inter-subject community variability. Each panel in this figure corresponds to the correlation of one behavioral measure for a single principal component.

**FIG. S3.**
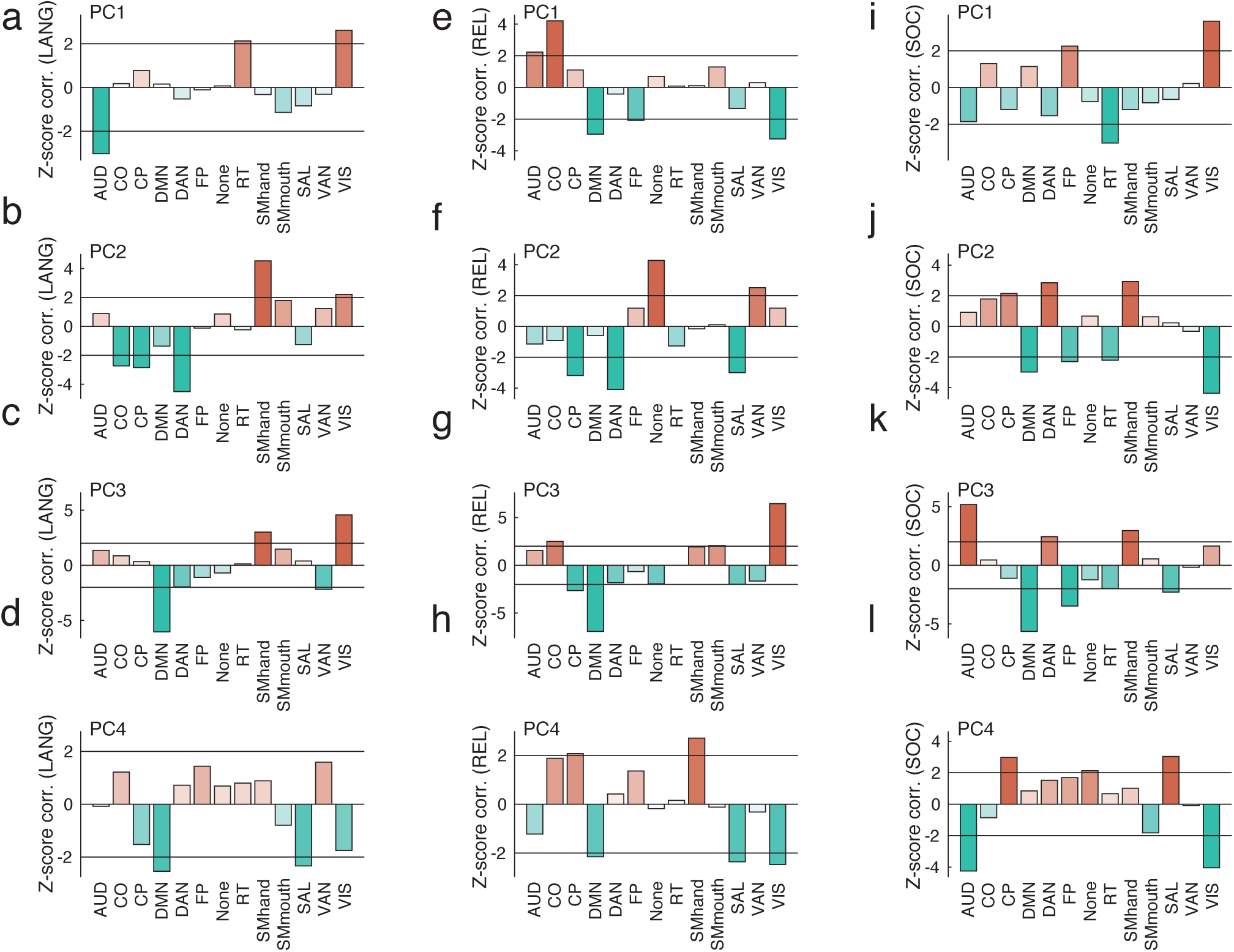
System-level maps depicting the correlation of community structure with measures of task performance. In this figure, we aggregated the regional correlation coefficients by cognitive systems and compared the mean system correlation to what we would expect by chance. Here, we report *z*-scored correlations. In general, larger positive *z*-scores indicate stronger than expected system-level correlations.

## Footnotes

1 Group-level community labels can be obtained in a number of ways, including data-driven methods such as community detection [13] or system labels taken from canonical brain atlases [5, 14]

